# svCapture: Efficient and specific detection of very low frequency structural variant junctions by error-minimized capture sequencing

**DOI:** 10.1101/2022.07.07.497948

**Authors:** Thomas E. Wilson, Samreen Ahmed, Jake Higgins, Jesse J. Salk, Thomas W. Glover

## Abstract

Error-corrected sequencing of genomic targets enriched by probe-based capture has become a standard approach for detecting single-nucleotide variants (SNVs) and small insertion/deletions (indels) present at very low variant allele frequencies. Less attention has been given to strategies for comparable detection of rare structural variant (SV) junctions, where different error mechanisms must be addressed. Working from cell samples with known SV properties, we demonstrate that Duplex Sequencing (DuplexSeq), which demands confirmation of variants on both strands of a source DNA molecule, eliminates false SV junctions arising from chimeric PCR. DuplexSeq could not address frequent intermolecular ligation artifacts that arise during Y-adapter addition prior to strand denaturation without requiring multiple source molecules. In contrast, tagmentation libraries coupled with data filtering based on strand family size greatly reduced both artifact classes and enabled efficient and specific detection of even single-molecule SV junctions. The throughput of SV capture sequencing (svCapture) and the high base-level accuracy of DuplexSeq provided detailed views of the microhomology profile and limited occurrence of *de novo* SNVs near the junctions of hundreds of sub-clonal and newly created SVs, suggesting end joining as a predominant formation mechanism. The open source svCapture pipeline enables rare SV detection as a routine addition to SNVs/indels in properly prepared capture sequencing libraries.

## INTRODUCTION

Detecting sequence variants in next generation sequencing (NGS) libraries demands that the signal from true variant DNA molecules rises above the background signal from pre-analytical and computational process errors. Historically, confidence was achieved by sequencing samples with moderate to high variant allele frequencies (VAFs) and demanding independent detection of variants in multiple source DNA molecules (1). As researchers have increasingly sought to apply NGS to samples with much lower VAFs, focus has shifted toward reducing the sequencing error baseline to improve signal-to-noise ratios. Applications where it is essential to detect sequence changes with very low VAFs include characterization of heterogeneity in tumor samples (2), studies of the nature and occurrence of mosaicism and somatic mutations in tissues (3-5), detection of genetic alterations induced by genotoxicants (6,7), and assessment of the off-target effects of gene editing methods (8,9).

Error minimization in NGS requires a detailed understanding of the mechanisms that create those errors (Figure 1A, Table 1). Error minimization strategies can be divided into two categories. We define “error suppression” as preventing pre-analytical, *i*.*e*., wet lab error mechanisms from impacting sequenced libraries so that false variants are never detected. PCR-free libraries and treatments to remove base damage are example error suppression strategies that eliminate or reduce PCR and base-copying artifacts, respectively (10). “Error correction” refers to bioinformatics strategies, sometimes supported by input library designs, that allow errors present in sequenced libraries to be recognized and discarded.

**Figure 1.**
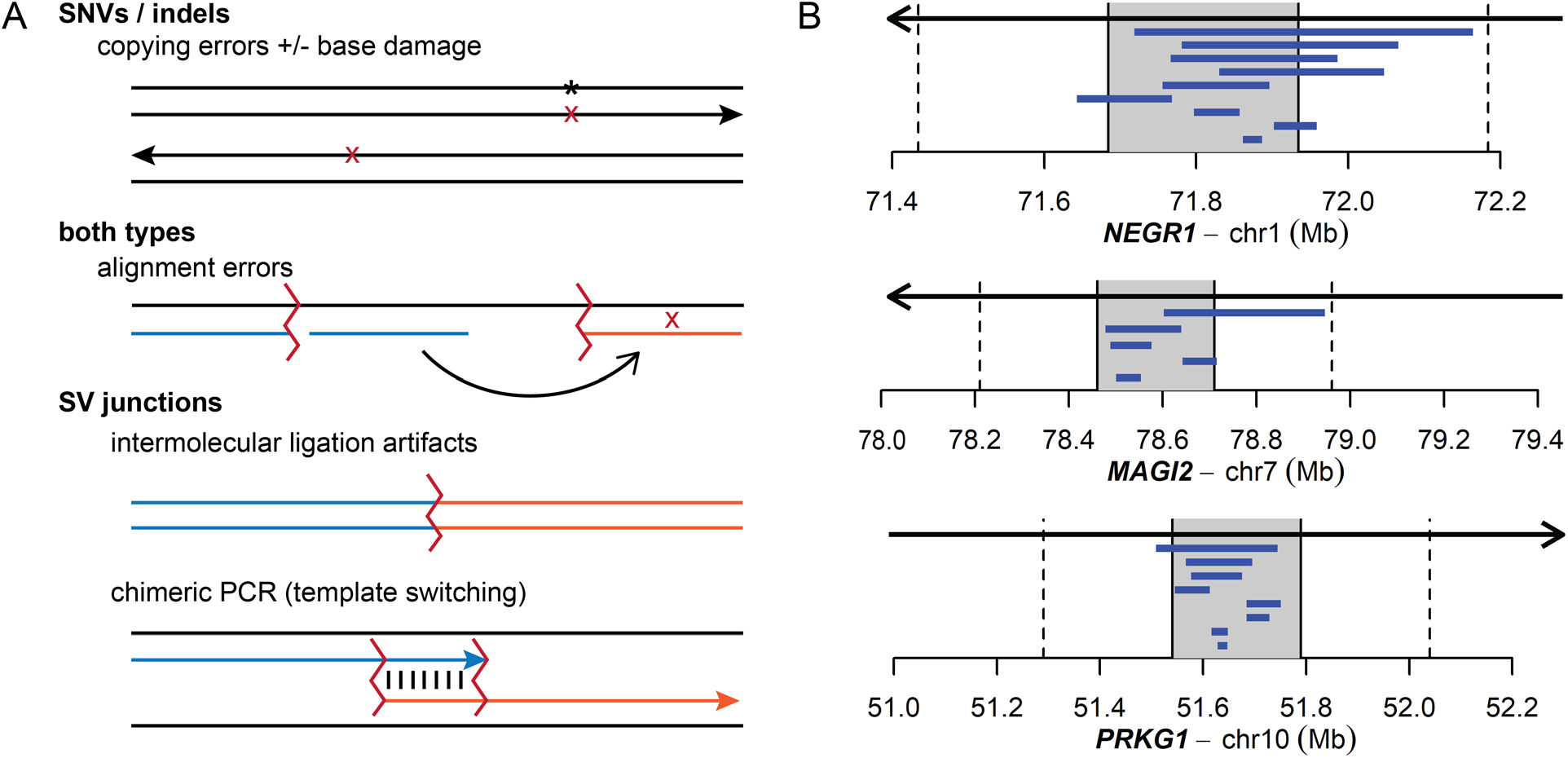
Design principles guiding svCapture and this study. **(A)** The most important error mechanisms giving rise to false calls for SNVs/indels and SVs. **(B)** The three targeted CFS genes. Arrows indicate the gene span and genome orientation. The central 250 kb shaded areas were targeted for capture (T regions). Vertical dashed lines mark the boundaries of the 250 kb adjacent (A) regions in triangle plots. Blue horizontal bars show the spans of overlapping SVs known to be present in sequenced clones based on prior microarray analysis.

**Table 1.** Properties of different classes of false and *bona fide* captured SV junctions.

| Parameter | Ligation Artifact | Chimeric PCR | True SV |
| --- | --- | --- | --- |
| Endpoint locations | Second endpoints randomly sampled from genome | SV endpoints paired within and between capture targets (but not adjacent regions) | SV endpoints paired within (but not between) capture targets and adjacent regions |
| Duplex molecules possible | Yes, artifact occurs before strand melting | No, artifact occurs after strand melting | Yes |
| Strand family sizes | Low to high | Low, arise late in PCR | Low to high |
| Endpoints shared with proper molecules | No | Yes, artifact templates pre-exist in library | No |
| Number of source DNA molecules | One | One | One to many, as VAF increases |
| Junction properties | Blunt ends or A/T insertions reflecting post-shearing A-tailing | Large microhomology to support high temperature annealing | Variable, predominantly 2-4 bp microhomology or small insertions |

Progress toward practical, error-minimized sequencing was advanced by the invention of Duplex Sequencing (DuplexSeq) (11,12). DuplexSeq is an error correction strategy in which unique molecular identifiers (UMIs) facilitate the recognition of the ends of individual source DNA molecules and the strands of those molecules that gave rise to sequenced read pairs. Computationally enforcing the detection of called variants on both DNA strands of a source molecule counteracts the most important error mechanisms giving rise to single-nucleotide (SNV) and small insertion/deletion variants (indels) – pre-existing base damage and polymerase copying errors (10) – which are inherently strand-specific (Figure 1A). DuplexSeq has been extended in derivative approaches dependent on the same core logic, such as NanoSeq and SaferSeqS (13,14). Together, duplex error-correction has enabled single-molecule SNV detection and is having a substantial impact on the applications above (15-18).

A limitation of most error correction strategies and studies is that they have not addressed the detection of rare structural variants (SVs). Like SNVs and indels, SVs are a critical form of mutagenesis associated with specific disease states and clastogenic genotoxicants (19-21). SVs alter large genome spans but their junctions are less frequent than SNVs and indels (22), increasing the need for stringency when assessing SV rates. Critically, the mechanisms that give rise to SV artifacts in sequencing libraries are very different from those that give rise to SNVs and indels, necessitating different error-minimization strategies (Figure 1A, Table 1). SV errors are created by three main mechanisms, listed here in their order of occurrence. Ligation artifacts arise from the joining of two source molecules by DNA ligase during adapter addition in libraries fragmented through sonication or non-tagmentation-based enzymatic fragmentation. Chimeric PCR occurs when a primer begins extension from one source DNA molecule only to switch to a different template in a subsequent cycle. Alignment errors occur when the aligner software reports an improper genomic location for a read or read segment.

Our efforts to address the challenges of error-minimized SV detection were motivated by our goal of defining the mechanisms of SV formation at common fragile sites (CFSs) (23-27). CFSs are genomic loci that are especially sensitive to replication stress and SV formation, representing genomic instability “hotspots” (28-30). Effective study of the molecular events occurring at CFSs demands that *de novo* SVs be detected as they form, when only a single source molecule will carry a given junction. Similar needs for ultra-rare SV detection exist in genotoxicant monitoring, characterization of low-level mosaicism in tissues, and other applications mentioned above. Here, we carefully analyze the frequency and nature of SV artifact classes in target capture libraries made from DNAs bearing SVs characterized *a priori* or increased population burdens of rare, induced SVs. DuplexSeq effectively addressed PCR artifact mechanisms, whereas tagmentation-based libraries coupled with bioinformatic filters addressed both PCR and ligation-mediated artifacts to yield signal-to-noise ratios suitable for monitoring rates of rare SV junction formation. The structures of hundreds of high confidence and accurately sequenced SV junctions suggests that they are most likely to arise by end joining.

## MATERIALS AND METHODS

### Human fibroblast cell line, clone mixtures, and stressed populations

Most samples analyzed in this study are derived from HF1, a TERT-immortalized primary human foreskin fibroblast cell line used in our prior SV studies (27,30). HF1 was propagated in Dulbecco’s Modified Eagle Medium supplemented with 13% fetal bovine serum, 4 mM L-Glutamine and 1× penicillin-streptomycin. Initial experiments used mixtures of previously described HF1 cell clones bearing known SVs (27,30). We recovered the clones from our frozen stocks, expanded them, and extracted DNA using the Qiagen Blood and Cell Culture Mini Kit (#13323). DNA was quantified using Qubit (ThermoFisher), diluted to 30ng/µl, and then mixed with DNAs from other clones or parental HF1 cells at known ratios. Further experiments used HF1 cell populations treated to induce *de novo* SVs. An exponentially dividing culture of HF1 cells was split and left untreated or exposed to either 0.2 µM or 0.6 µM aphidicolin (APH) in DMSO for 72 hours. Cells were allowed to recover without APH to complete SV junction formation for a further period of 24 hours. Cells were then split, seeded at 2×10^5^ cells, and expanded to confluence, after which DNA was extracted for sequencing.

### Target capture probe design and nomenclature

Capture probes were targeted to large genes according to the design parameters in Figure 1B and Table S1. Final probes were designed and synthesized by Twist Biosciences using their propriety algorithms to ensure specificity and quality and used as provided by the vendor. When denoting the correspondence of SV endpoints to target regions, “T” refers to bases within a capture target, “A” refers to bases in the 250 kb regions adjacent to each capture target, and “-” refers to all other genome bases. Uppercase letter pairs, *e*.*g*., “TA”, identify SVs with both ends in the same capture target, whereas lowercase letters, *e*.*g*., “tt”, identify all other unexpected, often artifactual, SVs.

### Duplex Sequencing libraries

DuplexSeq libraries were prepared by TwinStrand Biosciences Inc. (Seattle, WA) according to published procedures (12,17,18). Briefly, genomic DNA was fragmented by ultrasonication to a peak size of 300 bp, followed by end-repair, A-tailing, and DuplexSeq™ Adapter ligation. After indexing PCR, libraries were subjected to one round of overnight hybrid capture with biotinylated probes. Targets were purified with streptavidin magnetic beads, followed by washes and a final round of PCR prior to pooling and sequencing.

### Tagmentation sequencing libraries

Bead-based tagmentation libraries were prepared using the Illumina DNA Prep with enrichment, formerly called the Nextera Flex, kit (Illumina #20025523) by the University of Michigan Advanced Genomics Core. Tagmented libraries were prepared using an input of 300 ng of genomic DNA, IDT for Illumina unique-dual barcodes (Illumina #20027213), and library PCR amplification of 9 cycles. Libraries were quantified using Qubit and quality was checked using an Agilent Tapestation. Capture was then performed by pooling 500 ng of each library and hybridizing with 4 µl Twist Bioscience probes and 6 µl PCR grade water. Enrichment was completed with 12 cycles of PCR amplification.

## DNA sequencing

All sequencing reads were obtained in the 2 × 151 format using Illumina NovaSeq 6000 by the University of Michigan Advanced Genomics Core. Barcoded samples were pooled with each other and other users’ samples and subjected to a sequencing depth calculated to yield a projected coverage of ∼2,000-fold in the capture target regions based on prior experience. Library read counts, source DNA molecule yields, and enrichment values can be found in Table S2.

### svCapture data analysis tools and versions

Data analysis and visualization, including most data plots in figures, were accomplished using the svCapture pipeline and app in v1.0 of the svx-mdi-tools suite (https://github.com/wilsontelab/svx-mdi-tools),supported by genomicsmodulesin the main branch of the genomex-mdi-tools suite(https://github.com/wilsontelab/genomex-mdi-tools),asimplemented using the frameworksprovided bythe Michigan Data Interface (MDI,https://github.com/MiDataInt).Here, ‘pipeline’refers to high-performance computing actions executedusing variousprograms and custom shell, Perl, and Rscriptson the UniversityofMichigan GreatLakescluster, whereas’app’refersto RShiny graphical tools executed usingR 4.0.3 on a Windows 10 desktop computer. The job configuration scripts used tolaunchthe pipeline andassociated log filesthat list all options andprogram versionsare available at https://github.com/wilsontelab/publications/tree/main/svCapture-2022.

### svCapture pipeline

The svCapture pipeline’align’actionconverts putative DuplexSeqUMI sequences withinreadpairsto a UMI indexfrom 1 to 96, when applicable, allowing up toonemismatch betweentheobserved and expected UMIs. Reads withnon-matching UMIs are discarded. The fastp program (31)mergesoverlapping read pairs andenforcesread qualityfiltering. UMI indices and mergestates are recorded in read names.Thebwa memprogram(32,33)aligns readpairstotheGRCh38/hg38oranotherappropriate reference genome obtained fromIllumina iGenomeswithresults stored in name-sortedCRAM files.

The ‘collate’action performssource DNA molecule analysis. Twooutermost moleculepositionsareselectedfromaligned segmentspassing a map qualityfilter(MAPQ>20)toactas keysforcomparingreadpairsto each other.Read pairsthatsharethesame UMIindices, whenapplicable, andthe same endpointsand outer clip lengthswithin a 1 bp allowanceare groupedand referred to as a source DNA molecule.Forlibrariesusing ligated Y adapters, such asDuplexSeq, read pairswith opposite strandorientationsaregroupedaspartofthesamesourcemolecule. In contrast, differentread-pair orientationssharing the samemolecularendpoints mustarise from independentTn5 cleavageeventsandare considered differentsourcemolecules in), supported by genomics, supported by genomics modules in the main branch of the genomex-mdi-tools suite (https://github.com/wilsontelab/genomex-mdi-tools), as implemented using the frameworks provided by the Michigan Data Interface (MDI, https://github.com/MiDataInt). Here, “pipeline” refers to high-performance computing actions executed using various programs and custom shell, Perl, and R scripts on the University of Michigan Great Lakes cluster, whereas “app” refers to R Shiny graphical tools executed using R 4.0.3 on a Windows 10 desktop computer. The job configuration scripts used to launch the pipeline and associated log files that list all options and program versions are available at https://github.com/wilsontelab/publications/tree/main/svCapture-2022.

### svCapture pipeline

The svCapture pipeline “align” action converts putative DuplexSeq UMI sequences within read pairs to a UMI index from 1 to 96, when applicable, allowing up to one mismatch between the observed and expected UMIs. Reads with non-matching UMIs are discarded. The fastp program (31) merges overlapping read pairs and enforces read quality filtering. UMI indices and merge states are recorded in read names. The bwa mem program (32,33) aligns read pairs to the GRCh38/hg38 or another appropriate reference genome obtained from Illumina iGenomes with results stored in name-sorted CRAM files.

The “collate” action performs source DNA molecule analysis. Two outermost molecule positions are selected from aligned segments passing a map quality filter (MAPQ > 20) to act as keys for comparing read pairs to each other. Read pairs that share the same UMI indices, when applicable, and the same endpoints and outer clip lengths within a 1 bp allowance are grouped and referred to as a source DNA molecule. For libraries using ligated Y adapters, such as DuplexSeq, read pairs with opposite strand orientations are grouped as part of the same source molecule. In contrast, different read-pair orientations sharing the same molecular endpoints must arise from independent Tn5 cleavage events and are considered different source molecules in tagmentation libraries. Read-pair strand family sizes are counted, and a two-step consensus is constructed, first on each strand separately, and then, when available, between the two strands. We used a fractional threshold of 0.667 shared bases in at most 11 down-sampled read-pairs for consensus calling; sequence positions failing this criterion are masked as N bases. svCapture includes steps for alignment-guided merging of smaller read pair overlaps not handled by fastp. Consensus sequences are finally re-aligned to the reference genome.

The “extract” action examines final read alignments for patterns that predict an underlying SV junction (Figure S1). Source DNA molecules are described as a series of nodes at the endpoints of all alignment segments, each with a chromosome, position, and direction (left or right) that the molecule proceeds from that point with respect to the reference genome. Source DNA molecule replicates are defined by their shared outer nodes. Putative SV junction molecules are handled one at a time and characterized by their inner nodes. Node pairs defining junction edges are sorted to reflect the top strand of the reference genome in non-inverted segments in the order found in the source molecules. Associated alignment sequences and CIGAR strings are carried forward after reverse complementing the alignments within (but not flanking) inversion segments as defined relative to the conjoined reference genome, such that all sequences now represent contiguous reconstructed SV junctions. Nodes from proper and anomalous source molecules are compared to identify SV junctions that share endpoints with proper molecules.

The “find” action searches extracted node tables for sets of molecules with distinct outer node signatures but the same inner node signatures, *i*.*e*., independent source molecules crossing the same SV junction (Figure S1). Data are sorted by left nodes to allow molecules to be broken into sets where nodes are separated by less than the maximum proper insert size in the library as declared by bwa mem. Molecules within each set are re-sorted by the right node and the process repeated to yield an evidence set supporting each putative SV. Additional steps include further purging of likely duplicate source molecules with outer endpoints closer than 5 bp in total, the recovery of source DNA molecules that were clipped at the junction on their outer edges by at least 5 bases but not aligned, and the reconstruction of junction sequences lacking split read evidence by joining the inner clipped nodes of molecules where junctions fell in read gaps. A single source DNA molecule with the most central junction is chosen as a reference for parsing final junction descriptions, including SV type, correspondence to capture targets, and the nature of microhomologies and *de novo* insertions. All steps can be performed on a single sample or on a set of input samples analyzed together to ensure sensitive matching of even single source molecules between samples.

Finally, the pipeline “genotype” action compares the base content of SV calls to the non-SV DNA of the source individual. Proper molecules lacking SV junctions from CRAM files are analyzed by bcftools mpileup, call, and norm (34) to establish a VCF file with a biallelic genotype at every sufficiently covered genome position within and adjacent to each capture target region. The resulting unphased haplotypes are compared to consensus SV junction sequences obtained from all aligned segments over all source molecules in each SV set. SV consensus bases absent from either the reference genome or haplotype variant lists are marked as presumptive SNVs or indels created during SV junction formation.

### svCapture app and related downstream analysis

The svCapture app supports interactive visualization of SVs and the enforcement of further SV filters, such as demanding duplex molecules, minimum strand family sizes, on-target vs. off-target SV endpoints, and minimum and maximum numbers of supporting molecules or co-analyzed samples sharing the same SV.

Matching of expected vs. known SVs was achieved by first computationally comparing SV endpoints within a 25 kb allowance, given the imprecision of microarray calls. All potentially matching SVs were then manually examined for best matches, most of which were unambiguous as discussed below. VAFs were calculated for each sequenced SV as the count of total supporting molecules, including outer clips, divided by the average target region coverage. Expected VAFs were one half the clone frequency because all SVs were known to be heterozygous.

To assess the non-randomness of the positions of *de novo* SNVs and indels, the number of variant-containing SV consensuses crossing informative positions at each distance from the junction is counted to establish the expected distance distribution of randomly selected bases from those SVs. The observed number of variants is randomly selected from this weighted distribution over 10,000 iterations. The p-value is the fraction of iterations whose median is less than the median of the observed variant distances. A significant result indicates that SNVs and indels are more likely to occur near the SV junction (Mann-Whitney and other tests are also available).

## RESULTS

### Design of cell mixtures and capture targets for svCapture validation

To help optimize capture sequencing for rare SV detection, we drew on established human HF1 fibroblast cell clones bearing known replication-stress-induced SVs as established by genomic microarrays (27,30). We constructed a first set of five sample mixtures that were prepared, sequenced, and analyzed together that bore varying levels of different HF1 SV clones from 1% to 30% (Tables S3 and S4). The composition of most mixtures was known to the data analyst whereas a last sample was initially blinded and included unknown clone percentages and a subset of clones absent from other samples. A second set of five samples contained populations of HF1 cells treated with low doses of aphidicolin (APH), a commonly used agent of replication stress that is highly effective at inducing *de novo* SVs (30,35). To further promote predictable SV patterns, we selected three genes from published HF1 SV data (27,30) as high value hotspots of replication-stress-induced SV junction formation: *NEGR1* (0.89 Mb, chr1), *MAGI2* (1.44 Mb, chr7), and *PRKG1* (1.31 Mb, chr10). Capture probes targeted a territory of 250kb around the center each hotspot gene (Figure 1B, Table S1), which bears the highest burden of induced SVs (30). This represented a substantial increase in target span as compared to typical prior DuplexSeq libraries (15-18). Most SVs induced in these large genes are known to be deletions (30), which provided a further specificity test for method validation.

DNAs from the above sample sets were subjected to target capture and, initially, sequencing by DuplexSeq. A subset of the same DNAs was later sequenced again using tagmentation. We wrote a data analysis pipeline in the Michigan Data Interface because prior pipelines were optimized to address duplex consensus making or SV detection but not both (12,17,36). Like prior pipelines, svCapture groups read pairs by source molecule and strand, creates single-strand and, when possible, duplex consensus sequences, and realigns those consensuses to the reference genome. Additional features include read pair merging to maximize alignment specificity, persistence of all source molecules, and coordination of strand tracking with SV junction calling.

### DuplexSeq error correction eliminates chimeric PCR artifacts

Our entry rationale was that DuplexSeq would correct for chimeric PCR artifacts in capture libraries, which our early efforts suggested was the major confounder to SV detection. PCR is the last step in most capture protocols to create sufficient DNA mass for sequencing. It is problematic because it can promote chimeric PCR that is highly likely to fuse two molecules enriched by capture, mimicking the SV junctions we wish to detect. Chimeric PCR occurs after source DNA molecules are melted (Figure 1A) and should be amenable to duplex correction as it is highly unlikely that the same chimeric PCR event would occur twice independently, once in each template strand orientation. Because we know the properties of our source samples and genomic loci well (27,30), we can assert that apparent translocations between capture target regions are nearly always artifacts. True SVs will be mainly deletions with one captured endpoint within a target region (T) and the second endpoint within or adjacent (A) to the same region. Second endpoints large distances from the target region are again mostly artifacts.

We used plot devices described in Figure S2 to reveal the frequency, location, and properties of SV junctions detected by svCapture. Triangle plots of the locations of high-MAPQ, single-sample DuplexSeq SVs demonstrate a high burden of artifacts, most of which were confined within the boundaries of the capture target regions (Figures 2A and S3A). Most of those SV artifacts, including inter-target translocations, had large stretches of junctional microhomology with a peak at ∼8 to 9 bp (Figures 3A and S3E). This combination of features matches the expectations of post-capture chimeric PCR (Table 1). Consistently, requiring that at least one duplex molecule supported an SV call eliminated this error class (Figures 2B and S3B). Notably, duplex filtering was not required to correct for false chimeric PCR SV calls. Examining SV strand family sizes, *i*.*e*., the number of replicate read pairs from each source molecule strand (Figure S1A), showed that inter-target translocation SVs of the “tt” class rarely showed more than a single supporting read pair (Figure S4). Moreover, presumptive chimeric PCR events often shared one or both outer endpoints with proper molecules from the same library, the likely templates for late-cycle chimeric PCR (Figure S5A). Enforcing strand family size (Figures 2C and S3C) or shared endpoint (Figure S5B) filters proved effective for correcting chimeric PCR artifacts without sacrificing >50% of source molecules that failed duplex detection (Table S2).

**Figure 2.**
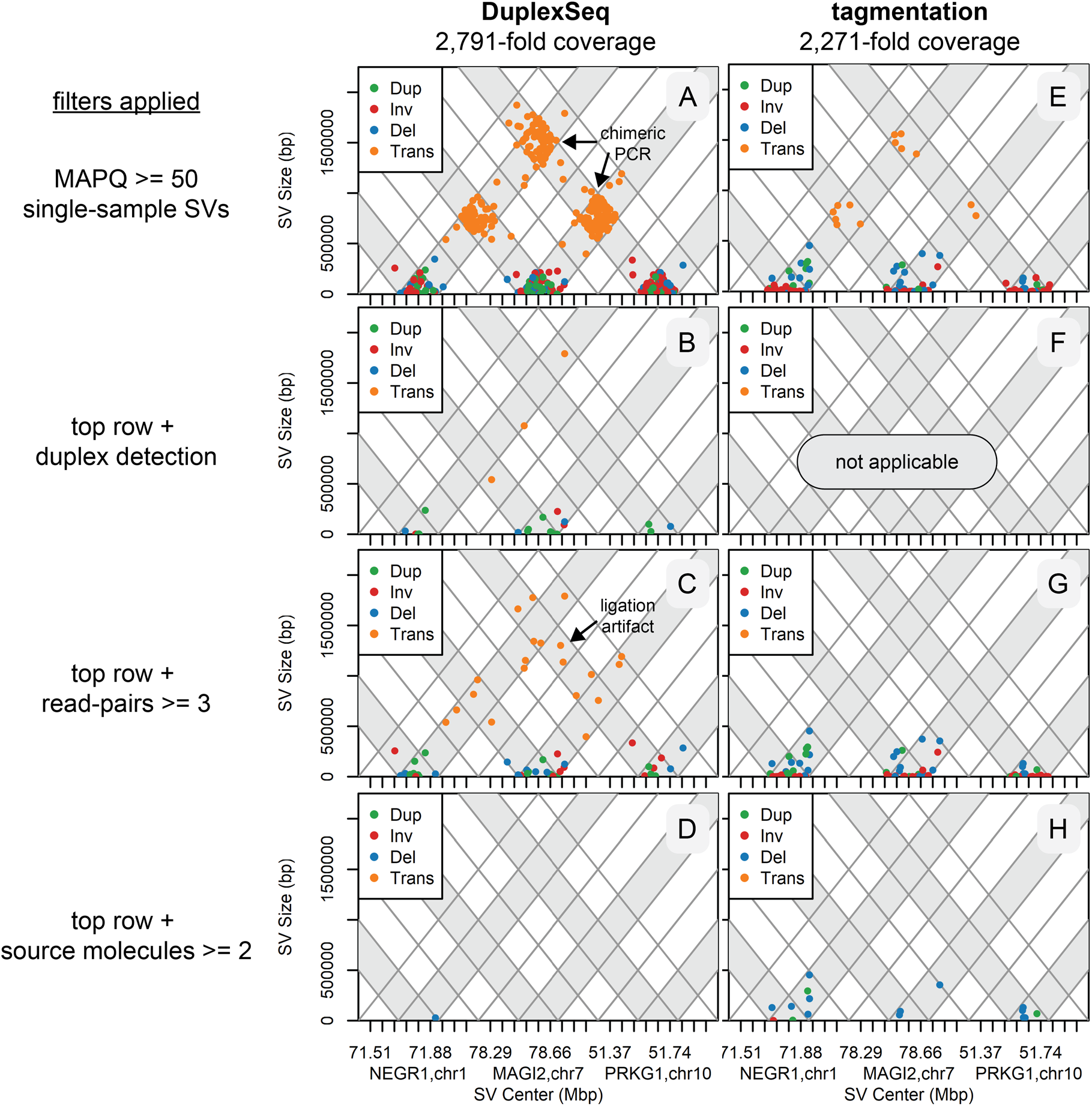
SV junction locations with and without error correction filters. Parallel triangle plots of untreated DuplexSeq and tagmentation libraries with similar target region coverage to minimize visualization bias. All plots are filtered for high-quality SV calls (MAPQ > 50 in each flanking alignment) detected in a single sample, the expected pattern for SV artifacts. **(A)** to **(D)** plot a single DuplexSeq library, whereas **(E)** to **(H)** plot two tagmentation libraries. **(A)** and **(E)** plot all detected SV junctions without further filtering. **(B)** and **(F)** further require at least one duplex molecule to plot the SV (tagmentation libraries do not support duplex assignments). **(C)** and **(G)** instead require that at least three read pairs (not source molecules) matched the SV. **(D)** and **(H)** instead require that at least two source DNA molecules matched the SV. See Figure S1 for definitions of read pairs and source molecules, Figure S2 for a guide to interpreting the plots, and Figure S3 for plots with all DuplexSeq data. Del, duplication; Inv, inversion; Del, deletion; Trans, translocation.

**Figure 3.**
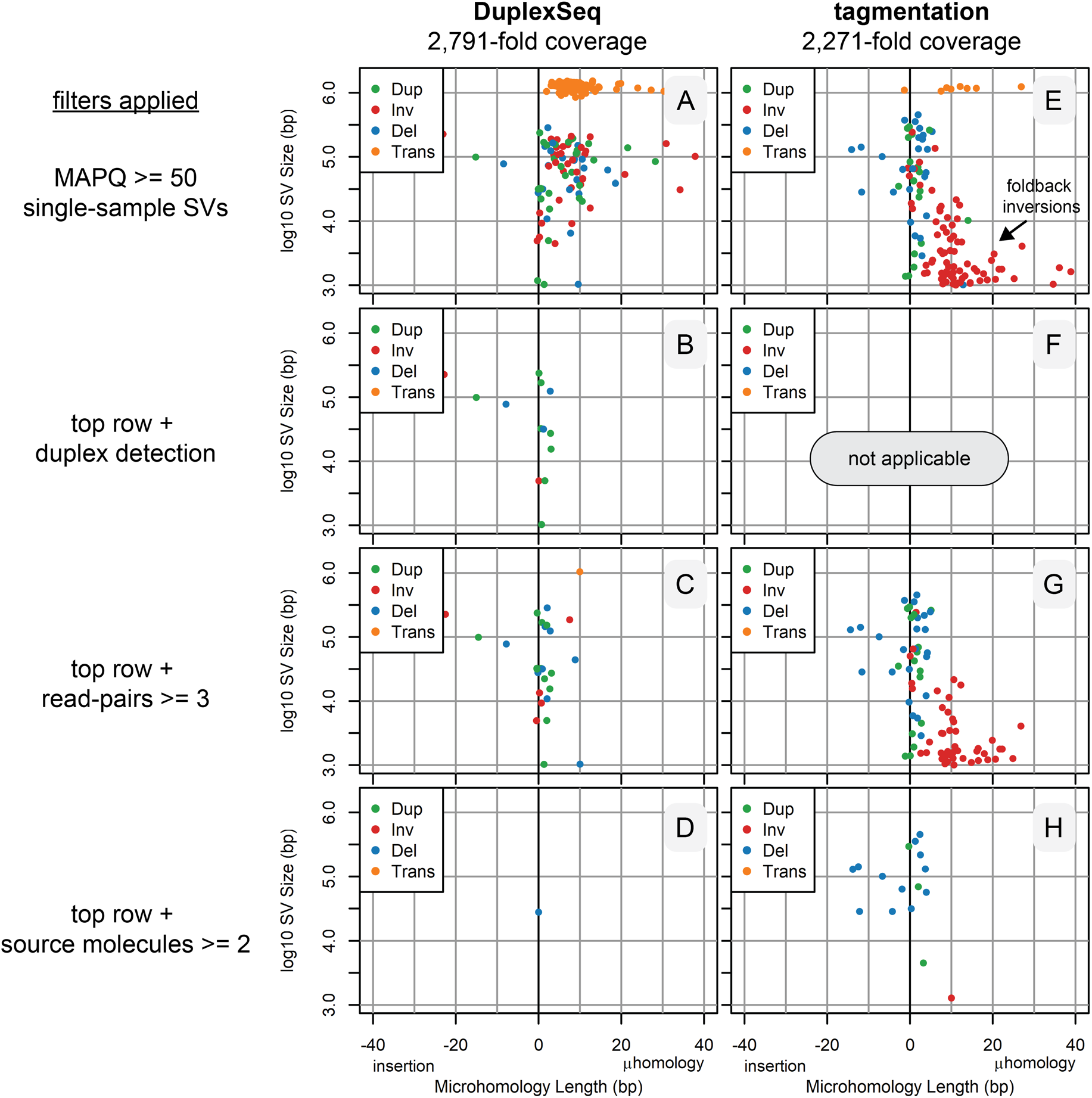
SV junction properties with and without error correction filters. Parallel junction property plots with the same samples and filters as Figure 2. See Figure 2 legend for details, Figure S2 for a guide to interpreting the plots, and Figure S3 for plots with all DuplexSeq data.

### DuplexSeq error correction alone cannot eliminate intermolecular ligation artifacts

Examining Figures 2C and S5B revealed a residual class of presumptive SV artifact junctions that did not respect the capture target boundaries, *e*.*g*., translocation junctions extending throughout the target (T) and adjacent (A) regions. Many of those remaining artifact junctions fell in the unsequenced gaps between read pairs (hence, they could not be plotted in Figure 3C), consistent with them being especially large molecules with central junctions, the expected pattern for intermolecular ligation artifacts. Consistently, this class of artifact junctions extended at the same baseline level throughout the untargeted genome (Figure 4A) and had a distinct junction profile characterized by blunt joints or single-base insertions or deletions (Figure 4B), as predicted for intermolecular ligation artifacts (1041 of 1293, 81%, of the 1-base insertions were A or T, suggestive of A-tailing). Because such artifacts arise prior to strand melting (Figure 1A), duplex filters alone are not expected to correct them (Table 1) and indeed they did not (Figures 2B, S3B, and 4C). Even with duplex filtering, tens of thousands of target-to-genome SV calls with predominantly blunt end junctions persisted that would confound many applications. Unsurprisingly, both ligation and chimeric PCR artifacts could be corrected by demanding multiple independent source molecules (Figure 2D), but that restriction prevents the detection of many rare, true SVs (see below).

**Figure 4.**
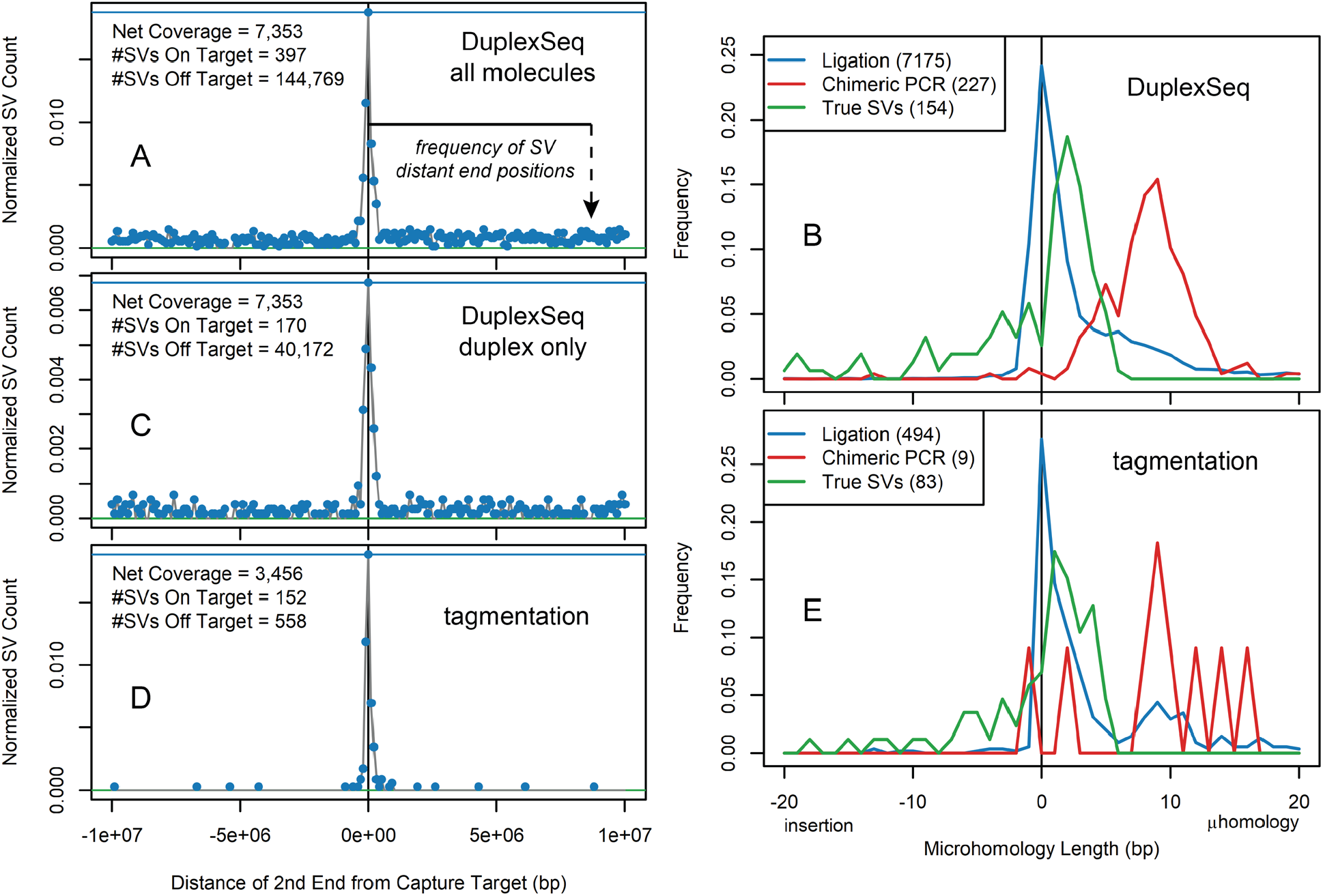
Intermolecular ligation is not corrected by DuplexSeq but is suppressed by tagmentation. All data represent single-sample, single-molecule, SVs >2 kb with a minimum MAPQ value of 50 for both SV alignments. **(A)** SVs from APH-treated DuplexSeq samples with at least 3 read pairs (to remove chimeric PCR events) in a 10 Mb window surrounding the capture target regions. For all plotted SVs, one end was within a capture target. The frequency of the other, more target-distant, junction positions are plotted per 100 kb bin. The central peak reflects APH-induced SVs in target and adjacent regions, whereas the baseline throughout the chromosomes reflects ligation artifacts. **(B)** Distribution of microhomology and *de novo* insertion lengths of three types of filtered junctions from DuplexSeq libraries. Presumed ligation artifacts are from (A), presumed chimeric PCR events are the tt-class translocation junctions filtered away from Figure 2A in Figure 2C, presumed true SVs are APH-induced deletion SVs of the TT or TA class with at least 3 read pairs. **(C)** Like (A), now also requiring at least one duplex molecule to plot the SV. The small apparent baseline reduction results from the loss of non-duplex molecules, not an increased SV call specificity. **(D)** Like (A), for all combined tagmentation libraries. Note the much lower rate of presumed ligation artifacts connecting the target regions to the rest of the genome. **(E)** Like (B), for the tagmentation libraries. The erratic chimeric PCR distribution results from the small number of such SVs.

### Tagmentation capture libraries largely, but incompletely, suppress ligation artifacts

Because there was no practical method for correcting ligation artifacts, we turned to error suppression to achieve this important goal. Because tagmentation libraries use the Tn5 transposase to connect adapters to genomic DNA molecules, not DNA ligase, ligation artifacts are not expected (Table 1). Comparing tagmentation to DuplexSeq libraries in the absence of any filters first revealed a new class of frequent, small inversions with large microhomologies that appeared uniquely in the tagmentation libraries (compare Figures 3A and 3E). Others have described that such errors arise by intramolecular foldback synthesis that is especially prevalent in tagmentation libraries, most likely during the extension of tagmented ends (37,38). Somewhat surprisingly, chimeric PCR artifact rates were lower in tagmentation libraries (compare Figures 2A and 2E), perhaps secondary to a greater capture target enrichment (Table S2). Family size filters were again highly effective at removing inter-target chimeric PCR artifacts as seen for DuplexSeq (Figure 2G and 3G). Most importantly, the rates of intermolecular ligation evident as “t-” class molecules were markedly reduced in tagmentation libraries (Figure 4D). The few such molecules that did occur shared a junction microhomology profile characterized by predominantly blunt ends (Figure 4E). This unexpected result suggests a low degree of ligation activity in tagmentation libraries (see Discussion).

### svCapture faithfully reports SV junction locations

To ensure that error correction and suppression were not eliminating true SV molecules, we examined our samples with mixed SV-containing clones, requiring three read pairs to call an SV. Figures 5A and 5B show a clear match to the SVs known to exist from prior microarray analysis down to a 1% allele frequency (compare detected SVs to the black circles denoting expected SVs). One expected 25 kb duplication in the *NEGR1* gene did not have an explicitly matching call, but we observed a nearby 4.5 kb duplication (red circle in Figure 5B) with unambiguous molecular evidence (Figures 5C and S6A). We interpret this as an error in microarray assessment of this SV, a method with much less precision in endpoint assignments, especially for duplications. We similarly observed apparently undetected SVs when we examined our initially blinded sample (Figure 5D). While we cannot rule out that some SVs were missed by sequencing, we again noted the presence of other strongly evidenced SV junctions leading us to question the precision of the microarray calls. For example, HF1 clone Scr1-14A had two ∼75 kb deletion calls in *MAGI2* by microarray (Table S4), whereas sequencing established the existence of a single, overlapping, unambiguous 149 kb deletion in the pool (red circle in Figure 5D, Figure S6B). Other unexpected but strongly evidenced SVs were also noted at low VAFs, such as the deletion highlighted by a red circle in Figure 5A and depicted in Figure S6C with six supporting molecules.

**Figure 5.**
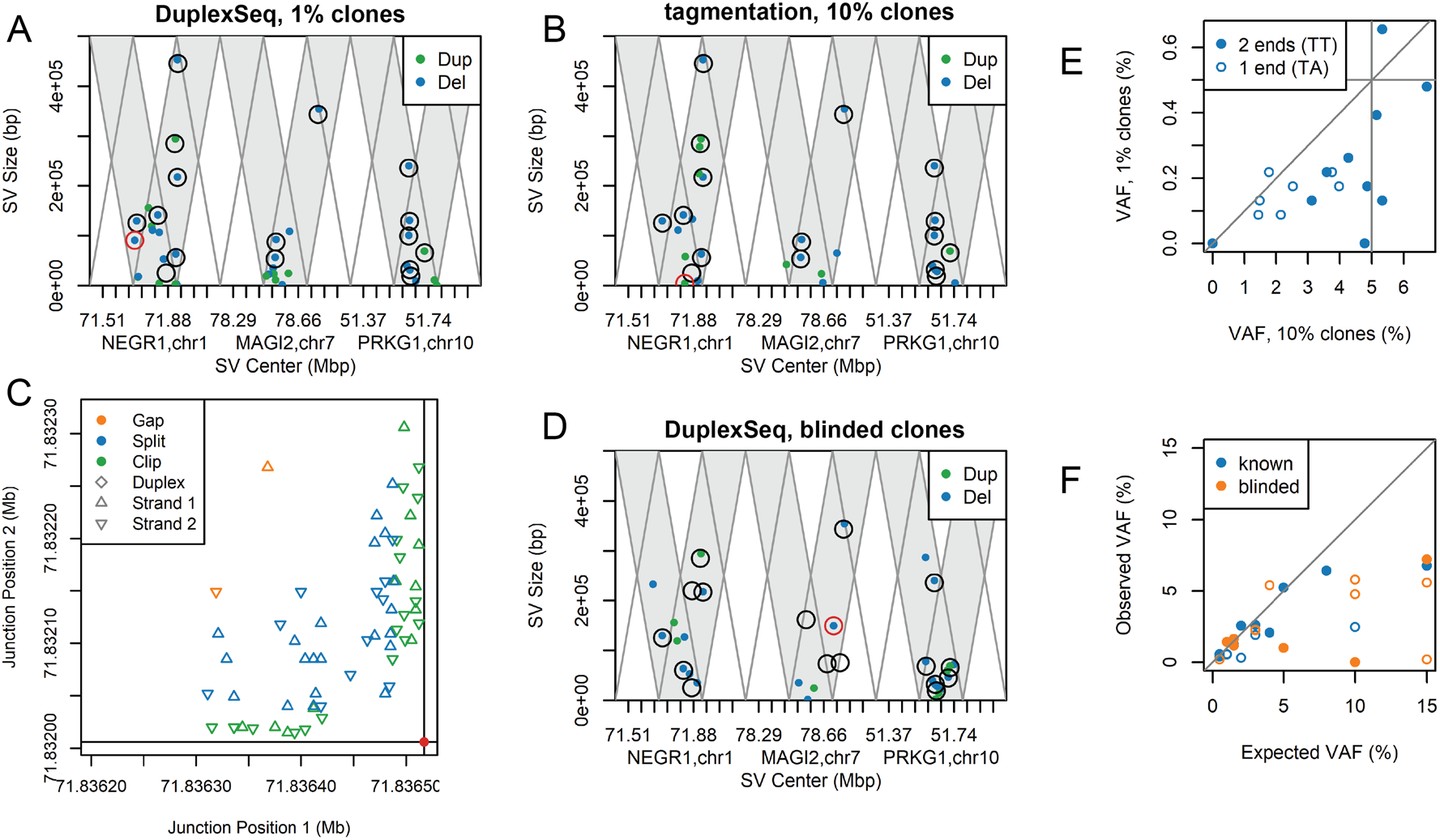
Sensitive and specific detection of known SVs down to 1% variant allele frequency. **(A)** and **(B)** Similar to Figure 2, showing duplications and deletions detected by at least three read pairs for the indicated HF1 mixed-clone samples. Black circles denote the positions of sub-clonal SVs expected based on prior microarray analysis. All but one matched both samples unambiguously. **(C)** Genome coordinates of the two outer positions of molecules supporting the SV highlighted by a red circle in panel (B), which likely accounts for the one call discrepancy in panels (A) and (B). The legend identifies the molecule types, which support a random distribution of multiple matching source molecules. **(D)** Like (A) and (B), for the blinded sample. See text for discussion of discrepancies and Figure S6B for a depiction of the SV marked with a red circle. **(E)** Correlation plot of the observed VAF for the same SVs from the 10% and 1% mixed-clone samples. Horizontal and vertical lines denote the expected values. Symbols indicate the number of SV ends that were inside a capture target region. **(F)** Correlation plot of the observed vs. expected VAF for the two indicated variable mixtures. Symbols are as in (E).

### svCapture reports SV allele frequencies with modest accuracy

To explore the quantitative efficiency and accuracy of svCapture, we plotted the observed vs. expected SV VAFs. While many SVs in the 10% mixture were recovered at approximately the expected VAF of 5%, those of the TA class showed lower VAFs (Figure 5E). SVs with only one end in a capture target region could thus be detected but at a lower efficiency, presumably due to reduced binding to capture probes. The 10% and 1% clone mixtures showed only a modest correlation (R = 0.74), in a pattern that initially suggested a reduced detection efficiency at lower VAF. However, plotting the observed vs. expected VAF for mixtures with variable content showed the opposite trend, with underrepresentation of the SVs with higher expected VAF (Figure 5F). SV recovery efficiency is thus variable and a function of several factors. Importantly, in this study one of those factors is the uncertain accuracy with which the clone mixtures were prepared.

### svCapture permits SV assessments at very low allele frequencies

While reliable detection of SV junctions at 1% VAF is valuable, it falls short of the needs of many applications. Therefore, we examined our APH-treated cell populations expected to carry a high burden of individually rare SVs across different cells. Figures 6A and 6B show the ready detection of increased SV junctions by either sonication-based or tagmentation-based svCapture following APH treatment of HF1 fibroblasts. In aggregate, these induced SVs were very often single-molecule detections (305 of 415, 73%), consistent with the expectations of *de novo* SV formation in a large cell population without clonal restriction. Induced SVs were strongly biased toward deletions (359 of 415, 87%), as we have previously observed at large gene loci (30), which, together with their induction by APH, provides excellent support for their existence in the source DNA. We further observed an apparent dose responsiveness of SV induction by APH and that tagmentation appeared more efficient than DuplexSeq at rare SV detection (Figure 6C), although more data are needed to draw firm conclusions on these points.

**Figure 6.**
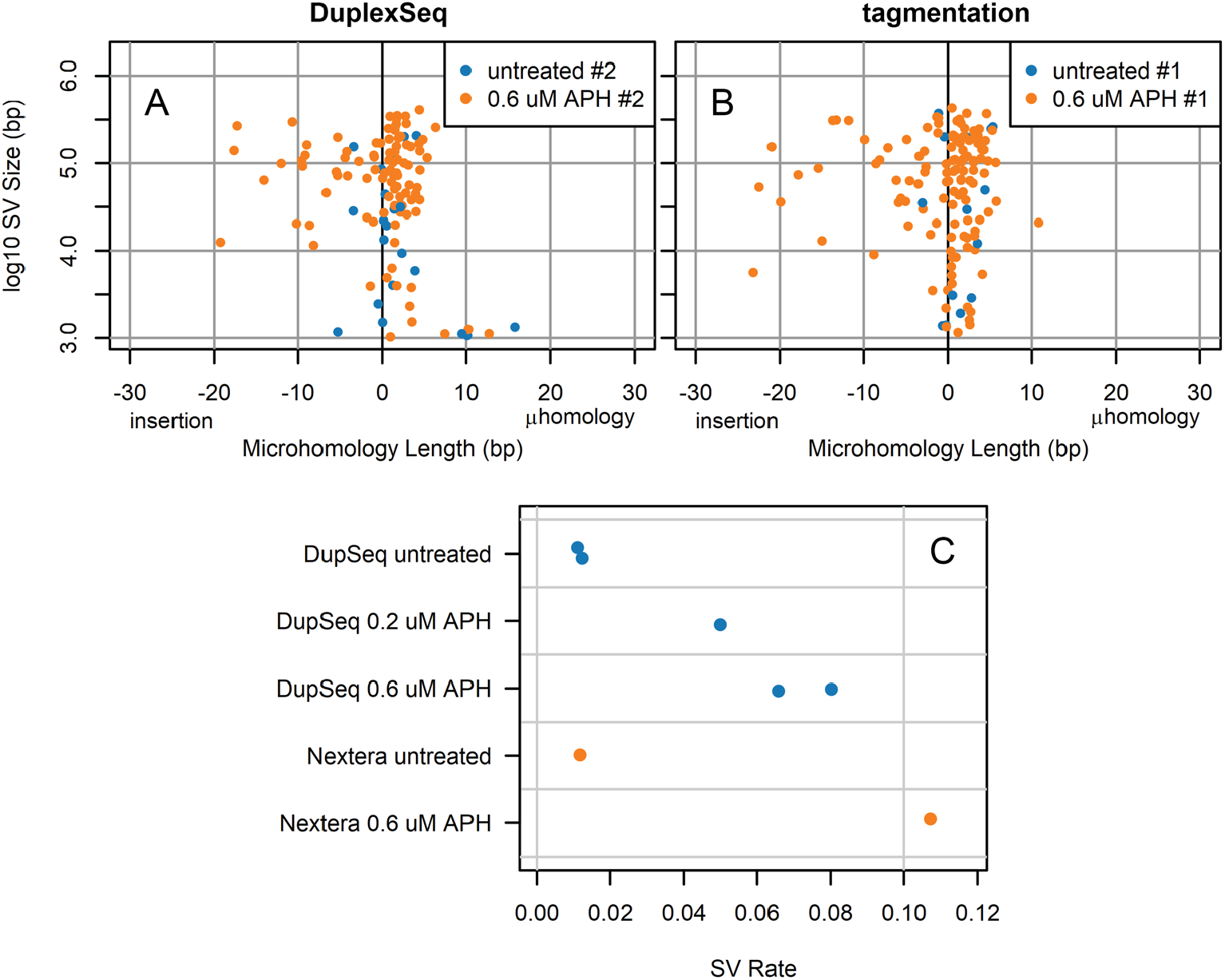
APH induces rare, detectable, often single-molecule deletion SVs in CFS genes. **(A)** and **(B)** Junction property plots comparing an untreated and APH-treated sample for each of the DuplexSeq and tagmentation-based svCapture approaches, showing consistent SV induction by APH. **(C)** SV formation rates, *i*.*e*., the number of observed junctions divided by the average on-target coverage, for all sequenced APH-treated sample libraries. Throughout, filters enforced a minimum MAPQ of 50 per alignment for SVs with at least 3 read pairs detected in a single co-analyzed sample.

### DuplexSeq reveals a low rate of *de novo* insertions, SNVs, and indels near SV junctions

We finally examined properties of SV junctions that are useful for inferring the associated DNA repair mechanisms. The hundreds of SVs induced by APH showed a junction profile distinct from either artifact class characterized by short, mainly 2 to 4 bp, microhomologies and a tail of junctions with *de novo* sequence insertions of up to 25 bp (Figures 4B, 4E and 6). These patterns provide further validation that these are valid SV calls distinct from the SV artifact classes. The high accuracy of DuplexSeq, coupled with abundant read depth for creating unphased haplotypes of the source individual, further allowed us to examine the rate of *de novo* SNVs and indels in the alignments flanking the SV junctions (see Figures 7A and S7 for an example). We measured a net rate of less than 1 in 1000 variant base positions in duplex SV calls that could not be accounted for by constitutional SNPs and indels in the HF1 genome (Figure 7B). The positions of the few variants observed in subclonal SVs from mixed clone samples appeared random, whereas there was a small but statistically significant bias (p <= 0.038) for the more numerous APH-induced SVs for being located closer to the junction than expected by random chance (Figure 7C).

**Figure 7.**
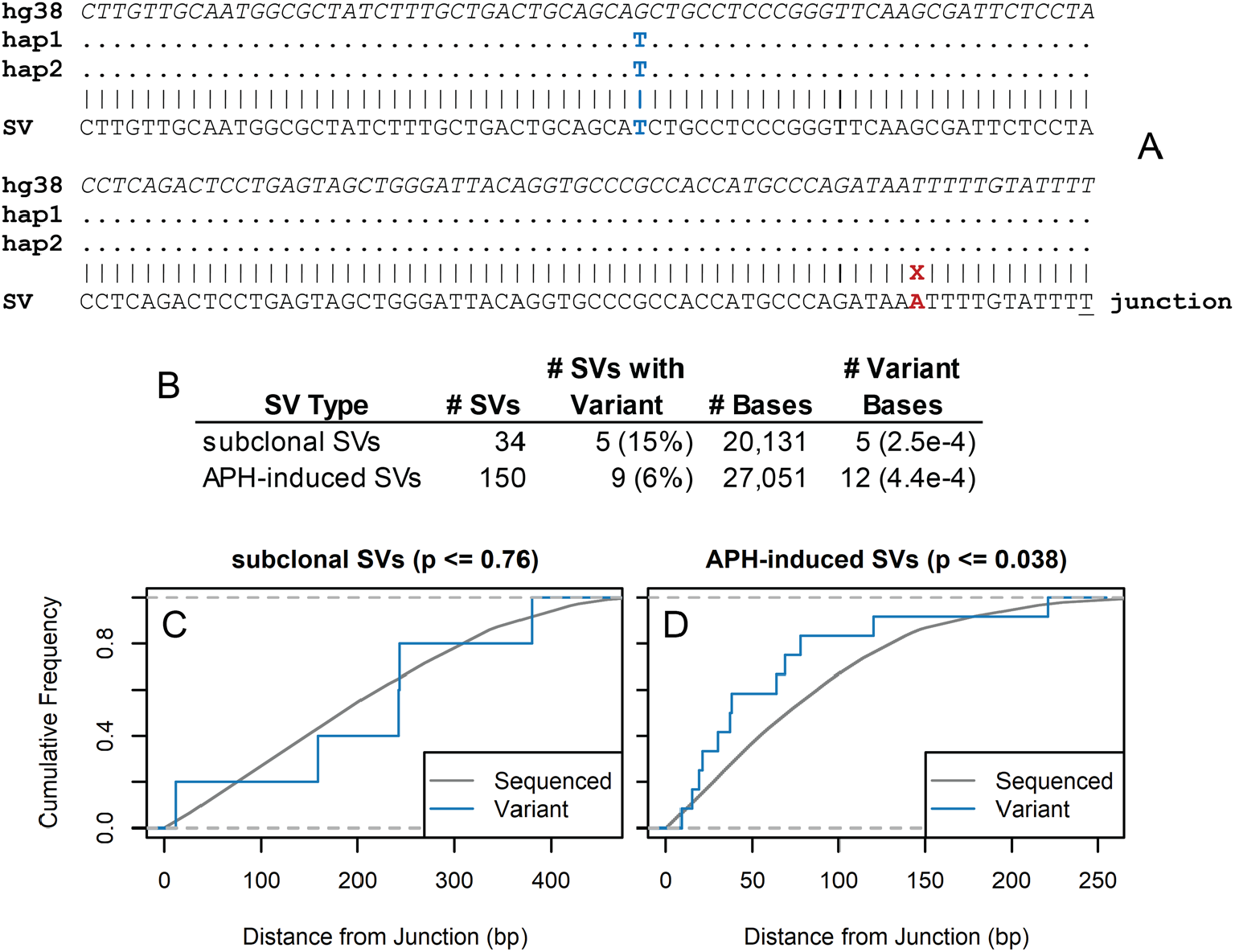
Low burden of *de novo* SNVs and indels near SV junctions. **(A)** An example SV sequence on the left side of the junction aligned to the reference genome (hg38) and the two identical haplotypes of the HF1 cell line (hap1 and hap2), showing a base match to a homozygous HF1 SNP (blue) and a *de novo* SNV in the SV located 11 bases internal to the 1-base microhomology (underlined). **(B)** Summary of the number and frequency of variant SVs and base positions for subclonal SVs present in two or more source molecules of the mixed clone DuplexSeq libraries, or single-sample SVs with 3 read pairs in the APH-treated DuplexSeq libraries. **(C)** and **(D)** Cumulative frequency plots of the number of sequenced bases (grey) and number of variant bases (blue) as a function of distance from the SV junction for the subclonal and APH-induced SVs, respectively.

## DISCUSSION

We describe svCapture, a powerful extension of error-minimized sequencing to SVs that will enable a variety of important downstream applications in molecular genetics, cancer biology, genetic toxicology, and other fields. A first conclusion from a rigorous analysis of multiple well-controlled samples is consistent with prior work with whole genome sequencing – that pre-analytical process errors, including intermolecular ligation and chimeric PCR, can be corrected by demanding that multiple independent source molecules match a junction (1,20). Demanding even two source molecules provides strong correction since all relevant wet lab process errors occur after fragmentation such that distinct molecule outer endpoints establish that an SV junction was present multiple times in the source DNA. Moreover, as compared to SNVs and indels, SV junctions provide abundant information to judge the specificity of SV matches between molecules. The processes for determining source molecule independence and matching SV junctions must be meticulous but are well described (20,36).

The challenge we addressed was extending SV detection to capture NGS where sequencing multiple independent source molecules is impractical or impossible due to low VAFs in input samples. Relevant applications include real-time monitoring of SV junction formation prior to replication in mechanistic studies, where only one source molecule could possibly carry the junction, or SVs arising just prior to the last mitotic division in neuronal or germ cell development (39-41) or during short-duration genetic toxicology studies (6,7,42). Even when SV junctions are sub-clonal, having been replicated during prior cell divisions, they are effectively unique at very low VAFs with respect to randomly sampled NGS libraries unless prohibitively deep sequencing is performed. At an average on-target coverage of 2,400 in bulk samples, we readily detected known SVs at 1% VAF with multiple molecules (Figures 5 and S6) but SVs would often yield single molecule detection at 0.1% VAF. Moreover, we observed only a modest ability of svCapture to accurately score VAFs. Multiple factors likely contribute to variable detection efficiency, including the number of ends captured, GC content, probe coverage, competition during capture and PCR, and others.

Of the three SV error classes (Figure 1A), we were the least concerned about alignment errors. Modern aligners, such as the bwa mem utility used in svCapture (32,33), provide reliable information regarding alignment confidence in the form MAPQ scores. Enforcing MAPQ filters reduces the detection of true SV junctions in repetitive DNA, but that lack of sensitivity is inherent to short-read libraries. Long-read technologies remain essential for characterizing SVs in repetitive regions (43-45). The greatest residual problem we have found regarding read alignment is for the subset of junctions containing short tandem repeats, where sequence changes arising during library preparation can cause aligners to give falsely high MAPQ scores.

In contrast, the remaining error classes – intermolecular ligation and chimeric PCR – readily yield false SV junctions with appropriately high MAPQ scores. Other methods are required to minimize their impact. Our goal was to improve SV signal-to-noise ratios by reducing artifact background, since true signals are obligatorily low in our target applications. Several computational filters could effectively correct for chimeric PCR errors, including duplex strand detection in DuplexSeq and high read-pair and low shared endpoint counts in all library types (Figures 2, 3, S3 and S5).

Thus, ligation artifacts proved of greatest concern. DuplexSeq logic could not correct for intermolecular ligation because the artifact occurs prior to strand melting and no other computational filter proved capable of doing so other than source molecule counts. However, libraries that use tagmentation to covalently bond adapter sequences to genomic DNA were highly effective at suppressing ligation artifacts from ever appearing in sequenced DNAs (Figure 4). Tagmented DNAs cannot typically support duplex error correction because they lack dual strand identification, but, as noted above, chimeric PCR can be corrected in other ways such that highly sensitive and specific detection of even single molecule SVs was achieved by tagmentation-based svCapture (Figures 2, 3, 5 and 6). Like others, we noted a class of small inversion artifacts in tagmentation libraries (37,38), but this does not impair the accurate calling of inversions larger than ∼1 kb or deletions, duplications, translocations.

Surprisingly, some apparent intermolecular ligation was observed in tagmentation libraries, even if greatly reduced relative to DuplexSeq. This inference is drawn because the artifacts in question had second SV ends spread evenly throughout the genome and the same predominantly blunt-end junction structure as seen with DuplexSeq (Figure 4). We do not know if the relevant phosphodiester bond formation is catalyzed by Tn5 itself or by some amount of ligase in the proprietary Illumina DNA Prep kits, although the former explanation seems most likely.

Importantly, any wet lab approach that prevents intermolecular ligation would suppress that artifact mechanism. The ideal would be a library prep that is both PCR-free and ligation-free. Indeed, PCR-free tagmentation libraries are now offered by vendors but with unspecified details as to how sequencer-ready libraries are created following Tn5 action. Quispe-Tintaya *et al*. reported Structural Variant Search (SVS) (42,46) for use on the Ion Torrent platform. Although SVS uses ligation to add second adapter strands it does so after homopolymer tailing by terminal deoxynucleotidyl transferase that effectively reduced intermolecular ligation. Importantly, these innovations in SV error suppression are largely restricted to whole genome sequencing, not target capture, and thus require a commitment to the higher costs associated with more sequencing as compared to known SV hotspots.

Data reported here represent the largest set of *de novo* SV junctions yet acquired in a controlled, prospective, experimental paradigm. They established in high accuracy calls two features of induced SV junctions, made possible in part by the application of DuplexSeq logic. First, the APH-induced junction sequence profile is characterized by short microhomologies and small *de novo* insertions (Figures 4 and 6) that are a strong match to DNA polymerase theta (POLQ) mediated end joining (TMEJ) (47,48). We also observed a low but measurable rate of *de novo* SNV acquisition near SV junctions that showed a biased localization in the first 50 bases on either side of the junction (Figure 7), consistent with a process confined to end sequences. This combination of features suggests that TMEJ or another end joining mechanism might create many SV junctions, with TMEJ favored since we previously showed that a canonical NHEJ factor, Xrcc4, is not required for SV junction formation (29). More data are needed, however, since both microhomologous junctions and error prone copying have also been invoked for replication-based SV formation models such as microhomology-mediated break-induced replication (49,50).

In summary, sensitive and specific detection of very rare SV junctions can be achieved in short-read capture libraries with sufficient attention to suppressing intermolecular ligation events by appropriate choices of library preparation protocols and correcting chimeric PCR artifacts by appropriate filters in data analysis pipelines. The svCapture procedure has a high degree of specificity and efficiency but modest quantitative accuracy for assessing VAFs. There are unavoidable limitations associated with short reads that include the inability to detect SV junctions in repetitive DNA or those mediated by recombination between longer regions of homology. Moreover, even in our optimized tagmentation approach the background of target-to-genome SV artifacts is not zero. Nevertheless, many important classes of nonhomologous SVs can be readily detected above very low background signals enabling inter-sample comparisons of the formation rates of even single-molecule SVs. The method is well suited to studies of SV formation mechanisms, somatic SVs, and SV-inducing genotoxicants.

## DATA AVAILABILITY

All code used in data analysis and figure generation are available for download via GitHub as part of the svx-mdi-tools repository at https://github.com/wilsontelab/svx-mdi-tools. HF1 cell svCapture sequencing data are available from the Database of Genotypes and Phenotypes (dbGaP) under accession PENDING.

## SUPPLEMENTARY DATA

Supplementary Data are available online.

## FUNDING

This work was supported by the National Institutes of Health [CA200731 to T.E.W and T.G.W., pilot project subaward of ES017885 to T.E.W]. Funding for open access charge: National Institutes of Health.

## CONFLICT OF INTEREST

The authors have no conflicts of interest regarding the contents of this manuscript.

## ACKNOWLEDGEMENTS

We thank Olivia Koues, Melissa Coon, and additional staff members at the Michigan Advanced Genomics Core for their expert assistance in library preparation and sequencing, Pam Bennett-Baker for support with library preparation and sequencing, Clint Valentine for helpful discussions regarding pipeline development, Anthony Nguyen for performing pilot studies of SNV formation in SV sequences, and Michael Hipp, Gabriel Pratt, Lindsey Williams, and Sharie Austin for analytical and project management support.

## Notes

https://github.com/wilsontelab/publications/tree/main/svCapture-2022

https://github.com/wilsontelab/svx-mdi-tools

### Competing Interest Statement

J.S. is Chief Executive Officer and Chief Scientific Officer of TwinStrand Biosciences, a company that sells Duplex Sequencing products for research uses. J.H. and J.S. are employees and equity holders at TwinStrand Biosciences Inc. and are authors on one or more duplex sequencing-related patents.

