## Supplemental Tables and Figures for "svCapture: Efficient and specific detection of very low frequency structural variant junctions by error-minimized capture sequencing"

### Table S1. Coordinates of target genes and capture regions

Chromosome coordinates are from the GRCh38/hg38 reference genome.

| Gene | Chrom | Gene Start | Gene End | Gene Size | Capture Start | Capture End | Capture Size |
|---|---|---|---|---|---|---|---|
| NEGR1 | chr1 | 71,395,943 | 72,282,539 | 887 kb | 71,684,317 | 71,934,317 | 250 kb |
| MAGI2 | chr7 | 78,017,055 | 79,453,667 | 1.44 Mb | 78,460,683 | 78,710,684 | 250 kb |
| PRKG1 | chr10 | 50,990,888 | 52,298,350 | 1.31 Mb | 51,540,240 | 51,790,240 | 250 kb |

### Table S2. Summary of library sequencing yields and capture enrichment

Yields are expressed as the number of sequenced read pairs and the number of source DNA molecules from which they were derived. Duplex rates are the fraction of on-target proper source molecules detected on both strands. Lines separate sets of libraries handled in the same batch.

| Method | Sample | Read Pairs | Molecules | On Target Coverage | Off Target Coverage | Enrichment | Duplex Rate |
|---|---|---|---|---|---|---|---|
| DuplexSeq | HF1 cell line | 292M | 69.9M | 2,447 | 5.63 | 435 | 48% |
| DuplexSeq | 10% clones | 265M | 64.9M | 2,363 | 5.28 | 448 | 48% |
| DuplexSeq | 1% clones | 263M | 62.7M | 2,290 | 4.69 | 489 | 47% |
| DuplexSeq | var. % clones | 323M | 71.5M | 2,260 | 5.50 | 411 | 48% |
| DuplexSeq | ? % clones | 294M | 69M | 1,969 | 5.17 | 381 | 48% |
| DuplexSeq | untreated, #1 | 314M | 81.7M | 2,791 | 7.23 | 386 | 48% |
| DuplexSeq | untreated, #2 | 289M | 92.3M | 2,669 | 7.80 | 342 | 49% |
| DuplexSeq | 0.2 uM APH, #1 | 337M | 98M | 2,682 | 8.15 | 329 | 49% |
| DuplexSeq | 0.6 uM APH, #1 | 348M | 94.1M | 2,344 | 9.05 | 259 | 48% |
| DuplexSeq | 0.6 uM APH, #2 | 272M | 80.4M | 2,327 | 7.07 | 329 | 47% |
| tagmentation | 10% clones | 113M | 6.8M | 1,083 | 0.11 | 9,669 | n.a. |
| tagmentation | untreated, #1 | 133M | 10.4M | 1,188 | 0.29 | 4,068 | n.a. |
| tagmentation | 0.6 uM APH, #2 | 120M | 7.3M | 1,186 | 0.12 | 9,561 | n.a. |

### Table S3. Composition of HF1 clone mixtures

| HF1 clone | HF1 cell line | 10% clones | 1% clones | var. % clones | ? % clones (blinded) |
|---|---|---|---|---|---|
| **GR36** | - | 10% | 1% | 1% | 3% |
| **Scr1-21A** | - | 10% | 1% | 2% | 6% |
| **GR68** | - | 10% | 1% | 3% | 20% |
| **O6A** | - | 10% | 1% | 4% | - |
| **Scr17A** | - | 10% | 1% | 6% | - |
| **GR8** | - | 10% | 1% | 8% | - |
| **E6A** | - | 10% | 1% | 10% | 2% |
| **Scr14A** | - | 10% | 1% | 16% | - |
| **Hp3-17** | - | 10% | 1% | 20% | 1% |
| **O3A** | - | 10% | 1% | 30% | - |
| **GR60** | - | - | - | - | 4% |
| **E8A** | - | - | - | - | 8% |
| **Scr1-2A** | - | - | - | - | 10% |
| **GR65** | - | - | - | - | 16% |
| **Scr1-14A** | - | - | - | - | 30% |
| **HF1 cell line** | 100% | - | 90% | - | - |

**Table S4. Prior coordinates of known SVs in clone mixtures**
Chromosome coordinates are from the GRCh38/hg38 reference genome, as determined by prior whole genomic microarray analysis.

| HF1 Clone | Chrom | CNV Start | CNV End | CNV Size |
|---|---|---|---|---|
| GR36 | chr10 | 51,616,453 | 51,648,536 | 32,083 |
| GR36 | chr10 | 51,628,463 | 51,647,510 | 19,047 |
| Scr1-21A | chr1 | 71,643,114 | 71,768,387 | 125,273 |
| Scr1-21A | chr1 | 71,830,439 | 72,047,181 | 216,742 |
| GR68 | chr10 | 51,508,820 | 51,744,414 | 235,594 |
| GR68 | chr7 | 78,602,605 | 78,946,165 | 343,560 |
| GR68 | chr1 | 71,861,295 | 71,886,554 | 25,259 |
| O6A | chr7 | 78,488,898 | 78,576,240 | 87,342 |
| O6A | chr1 | 71,718,794 | 72,164,112 | 445,318 |
| Scr17A | chr10 | 51,566,939 | 51,695,648 | 128,709 |
| Scr17A | chr1 | 71,902,277 | 71,958,434 | 56,157 |
| GR8 | chr10 | 51,576,665 | 51,676,014 | 99,349 |
| E6A | chr10 | 51,684,897 | 51,750,701 | 65,804 |
| Hp3-17 | chr1 | 71,781,124 | 72,066,242 | 285,118 |
| Scr14A | chr1 | 71,756,046 | 71,896,655 | 140,609 |
| O3A | chr7 | 78,500,709 | 78,554,235 | 53,526 |
| GR60 | chr1 | 71,307,910 | 71,420,871 | 112,961 |
| E8A | chr10 | 51,545,228 | 51,613,661 | 68,433 |
| Scr1-2A | chr1 | 71,797,112 | 71,857,252 | 60,140 |
| GR65 | chr10 | 45,876,166 | 45,892,175 | 16,009 |
| Scr1-14A | chr1 | 71,766,605 | 71,986,149 | 219,544 |
| Scr1-14A | chr10 | 51,684,414 | 51,729,219 | 44,805 |
| Scr1-14A | chr7 | 78,478,480 | 78,640,059 | 161,579 |
| Scr1-14A | chr7 | 78,642,674 | 78,715,891 | 73,217 |
| Scr1-14A | chr7 | 78,716,691 | 78,792,165 | 75,474 |

(figures begin on next page)

**Figure S1. Structural variant detection and molecule tracking in the svCapture pipeline**
**(A)** The relevant levels of sequencing redundancy. **(B)** The various ways that source DNA molecules (rows) might fall across a given SV junction (vertical red line) in both unmerged (top) and merged (bottom) read pairs. svCapture tracks the orientation of all alignments relative to anomalous splits, read-pair gaps, and inner and outer clips and correlates this information with the replicate, *i.e.*, consensus, detection of the sequenced strand(s) of each source molecule. Unique combinations of outer node pairs define source DNA molecules whereas unique combinations of inner node pairs define shared SV junctions. Depictions appear as the pipeline and app operate by considering the orientation of the DNA spans surrounding the SV junction as they were sequenced, not as they were aligned to the reference genome.

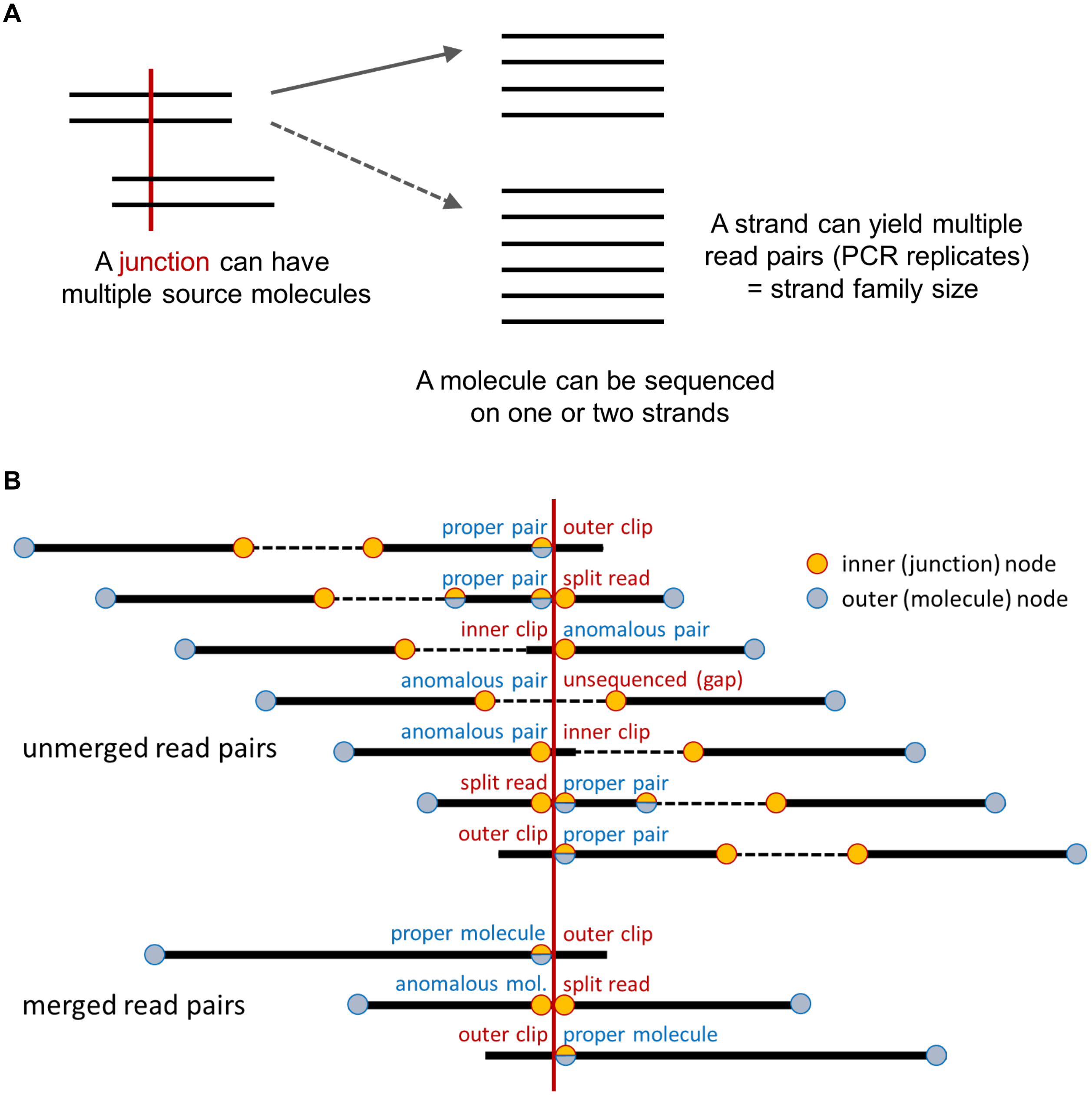

**Figure S2. Interpretive guide for plots**

**(A)** Triangular plots show SV endpoint locations relative to the capture target regions, which are highlighted by grey shading. The two endpoints of an SV are inferred by extrapolating diagonally to the x-axis and might be within or adjacent to a capture target or be a translocation between two different capture targets. **(B)** SV junctions are plotted to show the relationship of junction microhomology (x-axis) to SV size (y-axis). Translocations have no size and are plotted at ~1 Mb. Gaussian noise is applied to microhomology lengths to aid in visualization of all data points. Negative microhomology lengths denote *de novo* sequence insertions at junctions. Each dot corresponds to a single SV junction stratified by type (see color legend). Point plot order is randomized to minimize visualization bias due to overplotting.

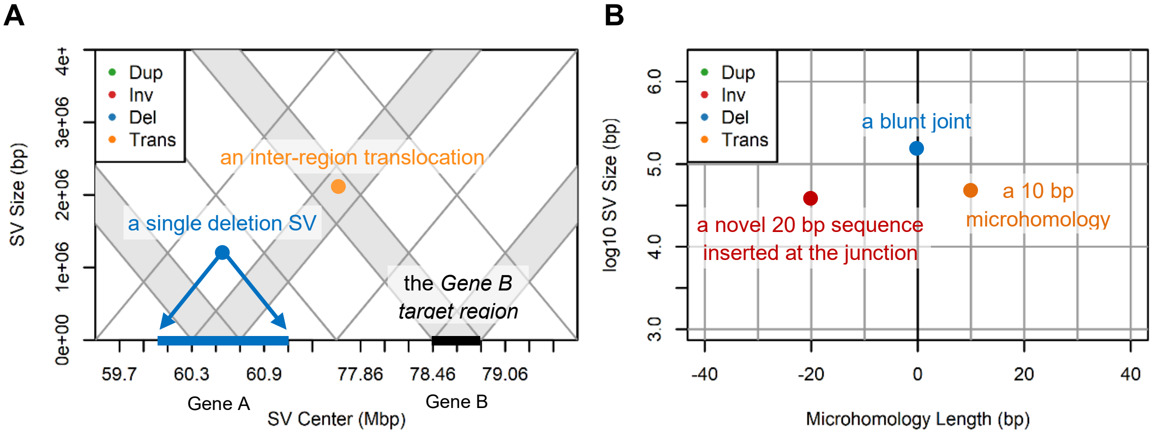

**Figure S3. Additional error correction plots from combined DuplexSeq samples**
Plots like Figures 2 and 3, now combining all untreated DuplexSeq samples at net 16,789-fold coverage for denser depictions. **(A)** to **(D)** are triangle plots of SV locations analogous to Figure 2A to 2D. **(E)** to **(H)** show SV properties analogous to Figure 3A to 3D.

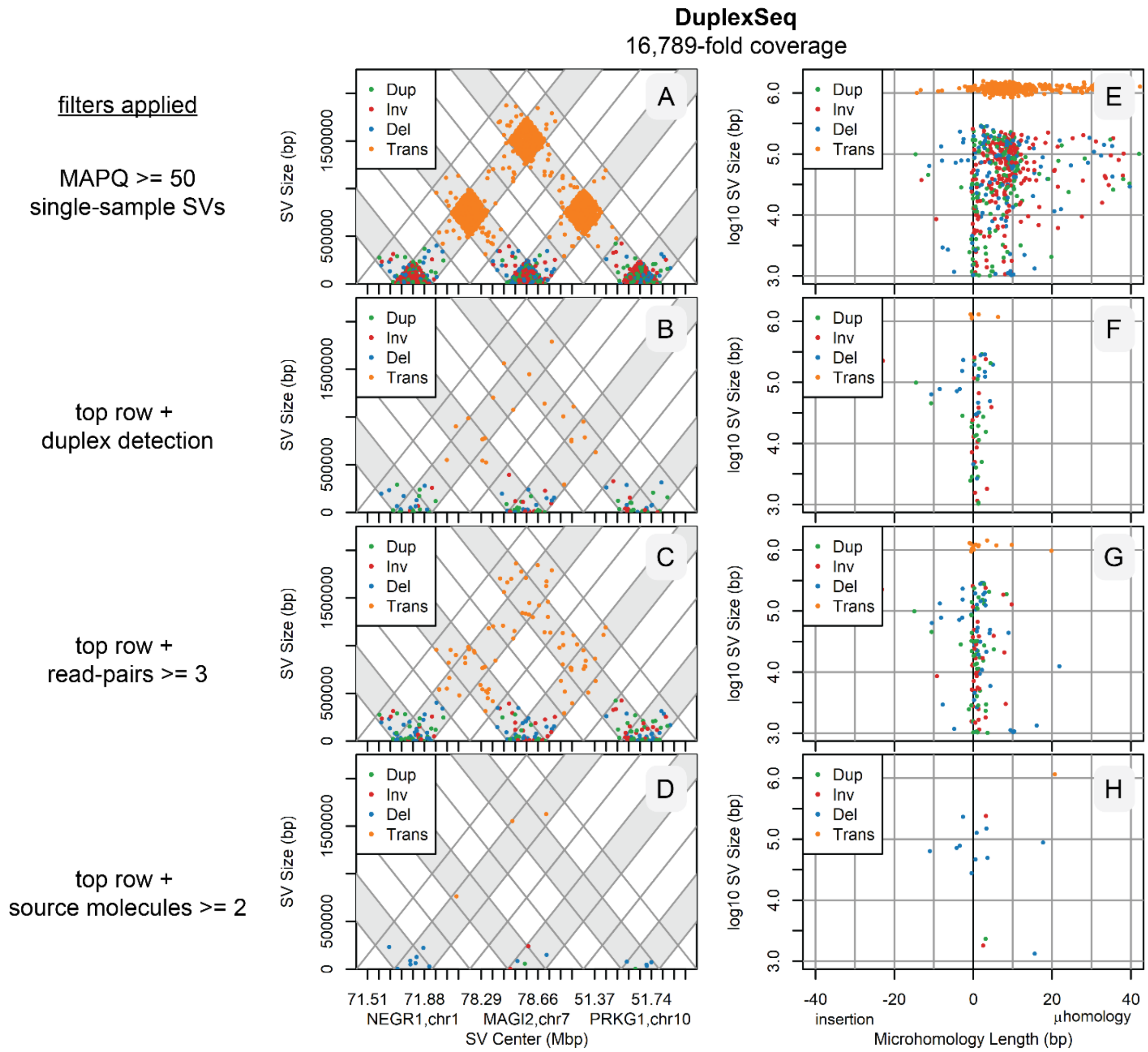

**Figure S4. Insert and strand family sizes for DuplexSeq and tagmentation libraries**

Figure panels show distributions of **(A)** and **(C)** library insert sizes and **(B)** and **(D)** source DNA molecule strand family sizes for representative **(A)** and **(B)** DuplexSeq and **(C)** and **(D)** Illumina DNA Prep tagmentation libraries. The same untreated HF1 cell input DNA was used for each. See Methods for interpreting “T” (target), “A” (adjacent), and “-“ (other) codes. In (C), the abundant class of “TT” proper molecules shows a 10 bp insert size periodicity reflecting the biophysical restrictions of bead-based tagmentation. Strand family size is the number of replicate read pairs corresponding to a given strand of a source DNA molecule. Solid lines are duplex molecules, dashed lines are single strand detections. For each method, “tt” translocation junctions (red) have very small strand family sizes, rarely more than a single read-pair.

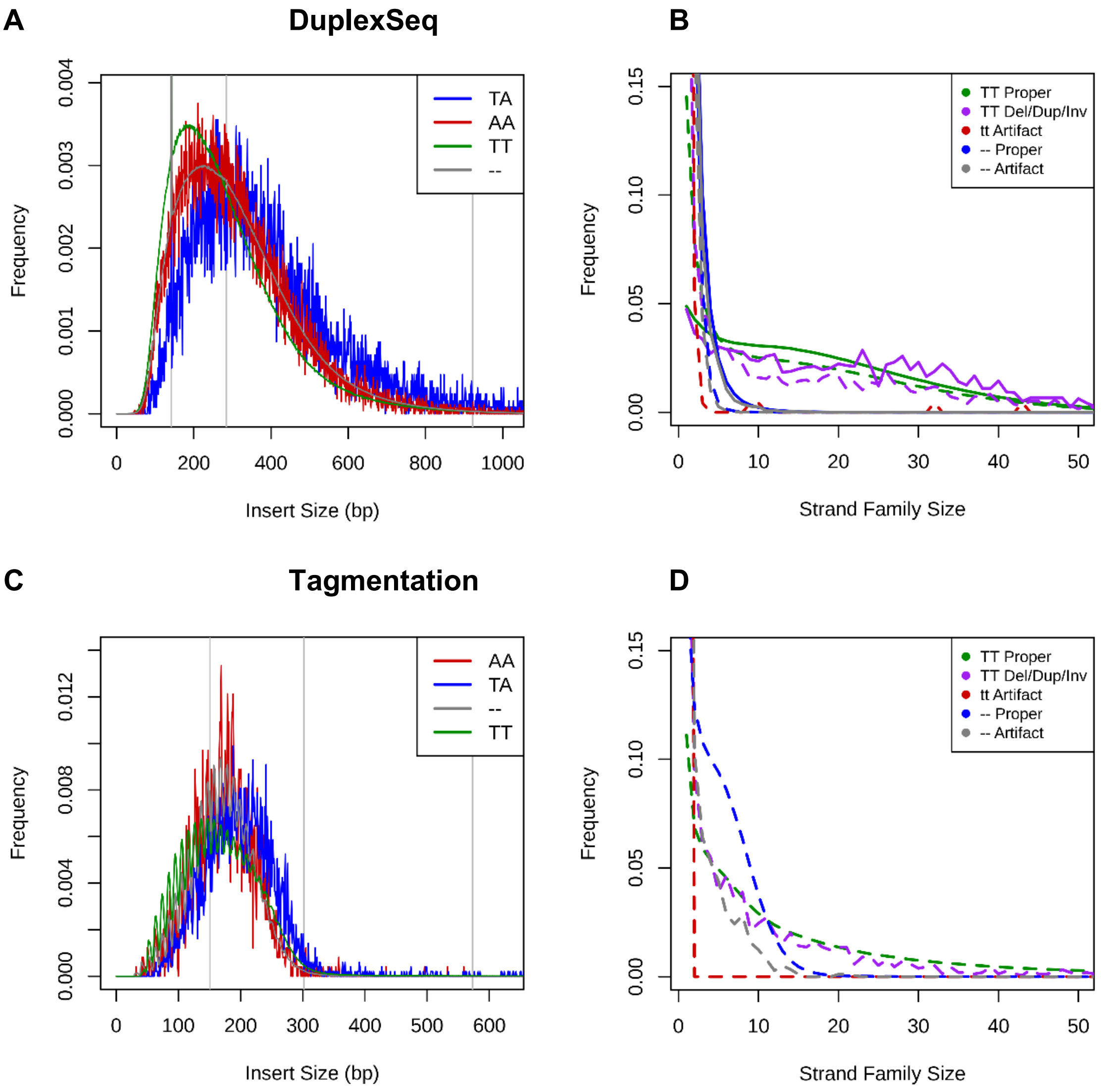

**Figure S5. DuplexSeq chimeric PCR artifacts often share endpoints with proper molecules (A)** Similar to Figure 3A to 3D, now plotting the fraction of fragment ends over all SV-supporting source DNA molecules that matched a proper source molecule in the same library. Most chimeric PCR artifacts at the peak of ~8 to 9 bp microhomologies shared both ends with proper molecules – the presumptive chimeric PCR templates – whereas the cluster of true SVs with a peak of 2 to 4 bp microhomologies did not. Data are from a pool of three DuplexSeq libraries from untreated HF1 cells. Gaussian noise was added to the y-axis values to aid in visualization of all data points, including single-molecule SVs. **(B)** Similar to Figure 2A to 2D for the same data as (A), now requiring that less than 25% of SV molecule ends be shared with a proper molecule.

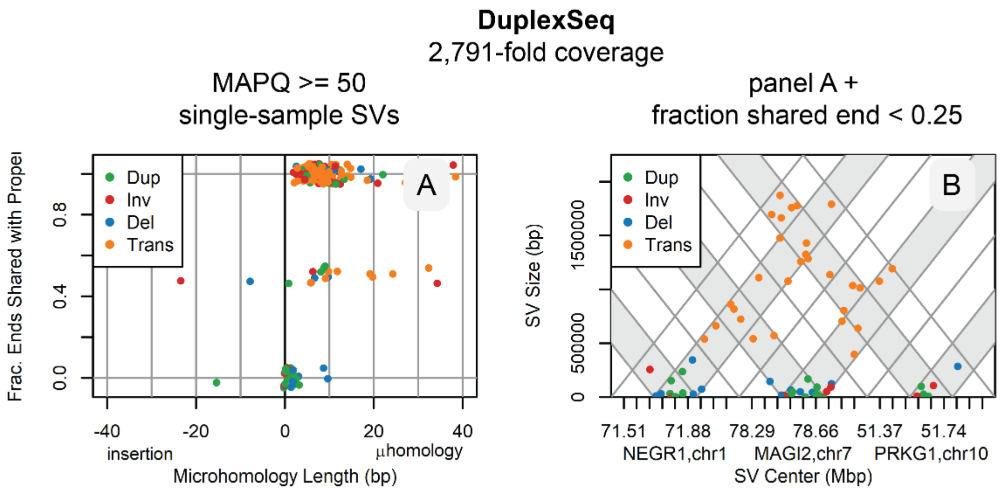

**Figure S6. Example molecule evidence for subclonal, mixed-clone SV junctions**

Drawings from the svCapture app depict all source molecules supporting example SV junctions. Colors reflect the base values at each position along the sequenced junction (A, green; C, blue; G, brown; T, red; N, black). The top half of each panel shows the alignments to the left side of the junction in bold colors, one alignment per row. The bottom half similarly shows the alignments to the right side of the junction. The clipped, *i.e.*, unaligned, portions are shown in faded colors. The position of the junction is indicated by vertical guidelines, with the internal yellow portions highlighting the microhomology span.

**A** – 4.5 kb duplication highlighted with a red circle in Figure 5B

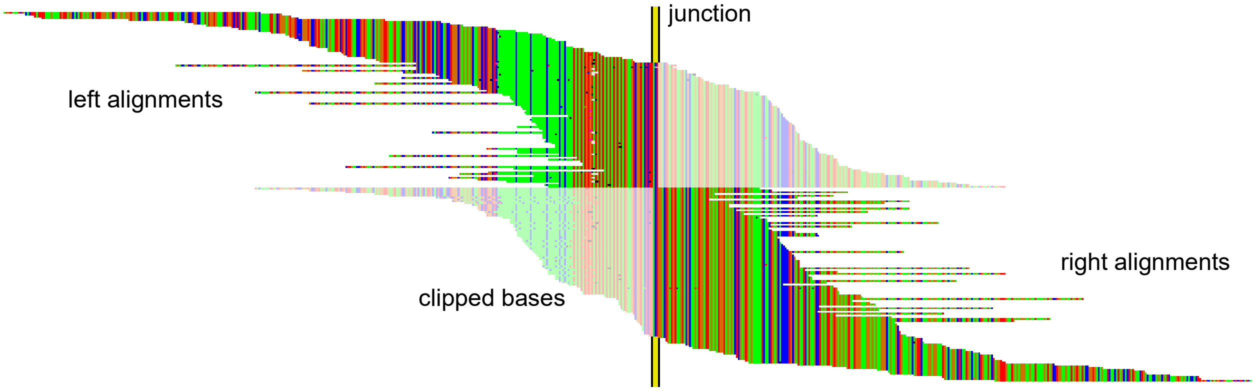


**B** – 149 kb deletion highlighted with a red circle in Figure 5D (outer edges cropped for clarity)

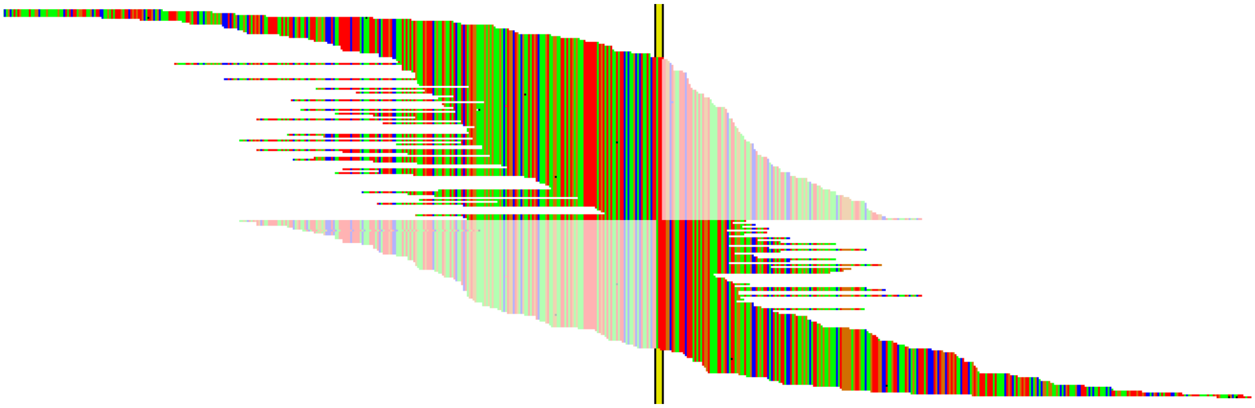

**C** – 91 kb deletion highlighted with a red circle in Figure 5A

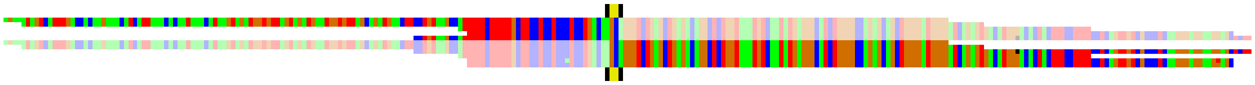

**Figure S7. Individual molecule evidence for a *de novo* SNV in an SV junction**

The same SV junction depicted in Figure 7A, now showing 10 of the 78 independent source molecule alignments (bottom lines) that sequenced across the variant base (red) as compared to the hg38 reference genome (top line). Lower case bases are the clipped bases in the split alignment. Only a portion of the total read sequences are shown.

```
CTACCTCAGACTCCTGAGTAGCTGGGATTACAGGTGCCCGCCACCATGCCCAGATAATTTTTGTATTTTTAGTAGAAACGGGGTTTCACCACGTTGG
|||||||||||||||||||||||||||||||||||||||||||||||||||||||||X|||||||||||
CTACCTCAGACTCCTGAGTAGCTGGGATTACAGGTGCCCGCCACCATGCCCAGATAAATTTTGTATTTTgaatagatgagatgtgtttttgctagct
CTACCTCAGACTCCTGAGTAGCTGGGATTACAGGTGCCCGCCACCATGCCCAGATAAATTTTGTATTTTgaatagatgagatgtgtttttgctagct
CTACCTCAGACTCCTGAGTAGCTGGGATTACAGGTGCCCGCCACCATGCCCAGATAAATTTTGTATTTTgaatagatgagatgtgtttttgctagct
CTACCTCAGACTCCTGAGTAGCTGGGATTACAGGTGCCCGCCACCATGCCCAGATAAATTTTGTATTTTgaatagatgagatgtgtttttgctagct
CTACCTCAGACTCCTGAGTAGCTGGGATTACAGGTGCCCGCCACCATGCCCAGATAAATTTTGTATTTTgaatagatgagatgtgtttttgctagct
CTACCTCAGACTCCTGAGTAGCTGGGATTACAGGTGCCCGCCACCATGCCCAGATAAATTTTGTATTTTgaatagatgagatgtgtttctgctagct
CTACCTCAGACTCCTGAGTAGCTGGGATTACAGGTGCCCGCCACCATGCCCAGATAAATTTTGTATTTTgaatagatgagatgtgtttttgctagct
CTACCTCAGACTCCTGAGTAGCTGGGATTACAGGTGCCCGCCACCATGCCCAGATAAATTTTGTATTTTgaatagatgagatgtgtttttgctagct
CTACCTCAGACTCCTGAGTAGCTGGGATTACAGGTGCCCGCCACCATGCCCAGATAAATTTTGTATTTTgaatagatgagatgtgtttttgctagct
CTACCTCAGACTCCTGAGTAGCTGGGATTACAGGTGCCCGCCACCATGCCCAGATAAATTTTGTATTTTgaatagatgagatgtgtttttgctagct
Etc.
```
